# Transcriptome guided metabolic network analysis reveals rearrangements of carbon flux distribution in *Neisseria gonorrhoeae* during neutrophil co-culture

**DOI:** 10.1101/2022.12.19.521143

**Authors:** Aimee D. Potter, Christopher M. Baiocco, Jason A. Papin, Alison K. Criss

## Abstract

The ability of bacterial pathogens to metabolically adapt to the environmental conditions of their hosts is critical to both colonization and invasive disease. Infection with *Neisseria gonorrhoeae* (the gonococcus, Gc) is characterized by the influx of neutrophils (PMNs), which fail to clear the bacteria and make antimicrobial products that can exacerbate tissue damage. The inability of the human host to clear Gc infection is particularly concerning in light of the emergence of strains that are resistant to all clinically recommended antibiotics. Bacterial metabolism represents a promising target for the development of new therapeutics against Gc. Here, we generated a curated genome-scale metabolic network reconstruction (GENRE) of Gc strain FA1090. This GENRE links genetic information to metabolic phenotypes and predicts Gc biomass synthesis and energy consumption. We validated this model with published data and in new results reported here. Contextualization of this model using the transcriptional profile of Gc exposed to PMNs revealed substantial rearrangements of Gc central metabolism and induction of Gc nutrient acquisition strategies for alternate carbon source use. These features enhanced the growth of Gc in the presence of neutrophils. From these results we conclude that the metabolic interplay between Gc and PMNs helps define infection outcomes. The use of transcriptional profiling and metabolic modeling to reveal new mechanisms by which Gc persists in the presence of PMNs uncovers unique aspects of metabolism in this fastidious bacterium, which could be targeted to block infection and thereby reduce the burden of gonorrhea in the human population.

**Importance:** The World Health Organization (WHO) designated *Neisseria gonorrhoeae* (Gc) as a high priority pathogen for research and development of new antimicrobials. Bacterial metabolism is a promising target for new antimicrobials, as metabolic enzymes are widely conserved among bacterial strains and are critical for nutrient acquisition and survival within the human host. Here we used genome-scale metabolic modeling to characterize the core metabolic pathways of this fastidious bacterium, and to uncover the pathways used by Gc during culture with primary human immune cells. These analyses revealed that Gc relies on different metabolic pathways during co-culture with human neutrophils than in rich media. Conditionally essential genes emerging from these analyses were validated experimentally. These results show that metabolic adaptation in the context of innate immunity is important to Gc pathogenesis. Identifying the metabolic pathways used by Gc during infection can highlight new therapeutic targets for drug-resistant gonorrhea.

## Introduction

*Neisseria gonorrhoeae* (the gonococcus, Gc) is the causative agent of the sexually transmitted infection gonorrhea. Gc is a human specific pathogen that is uniquely adapted to colonize human mucosal surfaces, where it survives despite initiating a robust inflammatory response and influx of innate immune cells, specifically polymorphonuclear leukocytes (PMNs, or neutrophils) (1). The mechanisms that Gc uses to resist PMN clearance remain incompletely understood. Gc encodes a relatively small repertoire of virulence factors compared to other pathogenic bacteria, and it has no known exotoxins (2). Instead, the success of Gc during human infection is related to its unique physiology, in particular its ability to exploit the resources in the host environment. Gc is a metabolic specialist that exhibits a limited carbon source preference, growing only on glucose, lactate, and pyruvate as sole carbon sources, suggesting that these nutrients are provided by the human host (3). As a human-adapted pathogen, many of the molecular determinants driving the specificity for the human host are required for nutrient acquisition (4). Metabolic gene products involved in lactate acquisition, nutrient metal import, and anaerobiosis are all required for full Gc virulence in models of infection ranging from cell culture to murine genital colonization to experimental human urethral challenge (5–12). However, many aspects of Gc metabolism remain undefined, such as the nutrients used by Gc in different infectious contexts and the core metabolic pathways required to sustain infection.

<u>Ge</u>nome-scale metabolic <u>n</u>etwork <u>re</u>constructions (GENREs) are a mathematical framework encompassing much of the known metabolic information on an organism (13). A draft GENRE can be generated with an annotated genome and several automated network reconstruction tools (14–17), then extensively manually curated using published literature and experimental data. GENREs can simulate all possible growth capabilities of an organism, which are then constrained by biological and physical parameters such as metabolite availability and optimized for a desired outcome, such as biomass production. GENREs enable large-scale, *in silico* manipulations of bacterial metabolism and have been used in a variety of applications including genome wide-knockout screens, synthetic lethal studies, and metabolic engineering that would otherwise be time-consuming and labor intensive to conduct (18). More recently, these tools have been used for the integration and interpretation of multi-omics data and applied to studies of human health and disease, including modeling of the metabolism of prominent human pathogens including *Mycobacterium tuberculosis, Staphylococcus aureus, Pseudomonas aeruginosa, Clostridioides difficile*, and *Salmonella typhimurium* (19–21). In contrast, there is no published model of Gc metabolism; while there is a GENRE for the related *N. meningitidis* (22), these two species are known to have key differences in their metabolism, for instance in sugar utilization (23). Moreover, few studies to date have applied metabolic modeling to pathogens in the context of immune cells, leaving a gap in knowledge of how immune-driven metabolic shifts shape bacterial metabolism. Systems-biology approaches are well suited to interrogating complex metabolic network interactions between organisms (24). These factors together make metabolic modeling an ideal platform for understanding novel metabolic drivers of Gc virulence.

Here, we present iNgo_557, a GENRE of Gc metabolism. This model enables the prediction of carbon source utilization and growth yields that recapitulate the behavior of Gc when grown in rich media. Metabolic network coverage in iNgo_557 includes genes, reactions, and metabolites that were initially identified by homology to a model of *N. meningitidis* and further curated using an automated model with support from literature evidence. The quality of iNgo_557 was further enhanced by update of standardized formatting and improving annotations. iNgo_557 was validated by comparing phenotypic predictions to experimental datasets and benchmarked with the MEMOTE test suite for assessing reconstruction quality. iNgo_557 was then contextualized with transcriptomic data that we generated for Gc grown with and without exposure to PMNs (Gene Expression Omnibus (GEO) database GSE123434), from which we identified and characterized unique metabolic features of the bacteria during an innate immune challenge. This GENRE of a clinically important, metabolically fastidious bacterium is a new resource for the *Neisseria* and microbial metabolic modeling communities. The insights into immune-driven metabolic shifts in Gc revealed by this transcriptionally-guided GENRE can inform the future development of therapeutic strategies to combat antibiotic-resistant gonorrhea.

## Results

### A genome-scale network reconstruction of *Neisseria gonorrhoeae* metabolism

We generated iNgo_557, a genome-scale metabolic network reconstruction of Gc strain FA1090, the type strain of Gc which is widely used and highly annotated. A published reconstruction of *N. meningitidis* M58 (Nmb_iTM560) served as the starting point (22) (**Fig 1A**). Nmb_iTM560 was based on the highly annotated iAF1260 reconstruction for *Escherichia coli* and was built using the Biochemical, Genetic and Genomic knowledge base (BIGG) framework (25). We identified homologous genes between *N. meningitidis* M58 (AE002098.2) and *N. gonorrhoeae* FA1090 (AE004969.1) using an homology matrix based workflow for generating high quality multi-strain genome-scale metabolic models (26). Gc and *N. meningitidis* were found to share significant homology across large stretches of the genome, particularly for metabolic genes (27): Of the 560 genes, 1519 reactions, and 1297 metabolites originally present in Nmb_iTM560, 494 genes, 1223 reactions, and 1189 metabolites were preserved in iNgo_557 based on homology (**Dataset S1**). Orphan reactions from Nmb_iTM560 with no corresponding gene were included in the initial Gc reconstruction and de-orphaned or removed where possible during manual curation. The format was updated to SBML Level 3, the most up-to-date community standard (28). Gene, reaction, and metabolite annotations were updated from KEGG, PATRIC, Uniprot, MetaNetX, MetaCyc, PubMLST, and BIGG databases wherever possible (25, 29–34).

**Fig 1.**
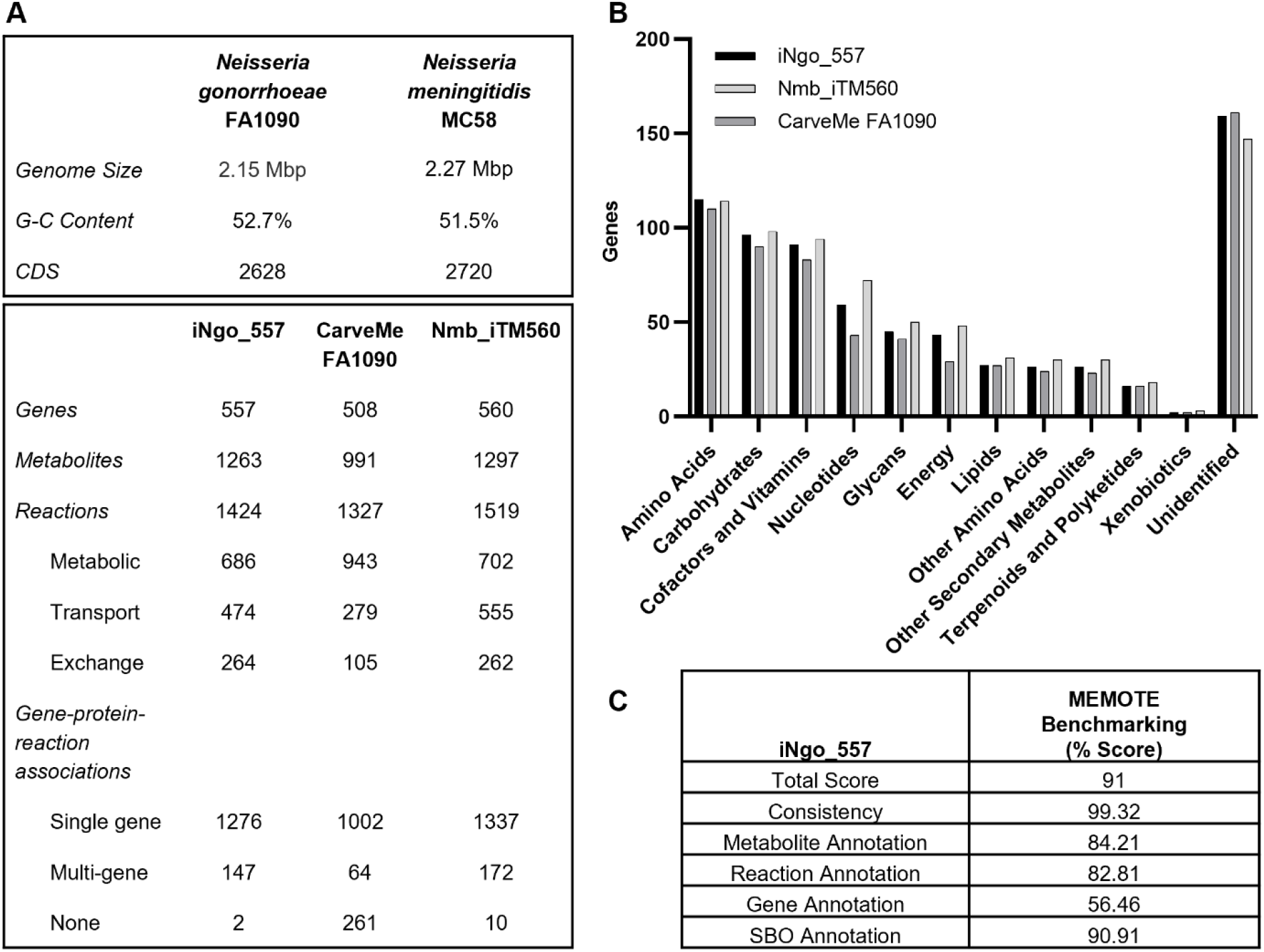
Genome-scale metabolic model of Gc strain FA1090. (A) (Upper panel) Comparison of Gc strain FA1090 and *N. meningitidis* strain MC58. (Lower panel) roperties of iNgo_557, CarveMe FA1090, and Nmb_iTM560. (B) Comparison of KEGG functional annotations for genes present in the three models. Some genes have multiple functions and are assigned to multiple categories. (C) MEMOTE benchmarking scores of iNgo_557.

Characteristics of the original Nmb_iTM560 model were conserved, including the presence of a periplasmic compartment, simplified cytochrome respiration pathways, iron acquisition pathways from ferric iron and host proteins, and a biomass equation that reflects the neisserial cell composition. Targets for manual curation of the automated reconstruction in iNgo_557 included complete resolution of mass and charge balance inconsistencies, the resolution of import and export loops, removal of carbohydrate import through the phosphotransferase system (which is not functional in pathogenic *Neisseria*) (23), addition of amino acid catabolism pathways, curation of lipooligosaccharide synthesis for Gc and its addition to the biomass equation, modification of the biomass composition for Gc where appropriate, and simplification of lipid biosynthesis (**Dataset S1**). Additionally, catalytic cofactors such as biotin, thiamine pyrophosphate, pyridoxal-5-phosphate, iron, zinc, manganese, NAD, and FAD were removed. In Nmb_iTM560, these cofactors were included as consumed reactants in reactions to reflect biological requirements for biomass production (35). While useful, the presence of these artificially consumed cofactors impedes the accurate representation of reaction stoichiometries in the reconstruction.

This homology-based reconstruction process was incapable of identifying Gc-specific genes that were not present in Nmb_iTM560 (26). Therefore, to expand the metabolic coverage of iNgo_557 for metabolic pathways that are unique to Gc, genes and their corresponding reactions/metabolites were added from an automated reconstruction in the BIGG namespace of Gc FA1090 that was generated using CarveMe (36). Each of the unique genes identified by CarveMe was manually evaluated. Of the 508 genes with metabolic functions predicted by CarveMe, 388 were already present in the model. CarveMe identified an additional 39 genes and corresponding reactions that were supported by manual evaluation of the literature, and they were subsequently included in iNgo_557 (**Fig 1A, Dataset S1**) (36). The remaining 81 genes identified by CarveMe did not have sufficient evidence to support the assigned metabolic function and were not added (**Dataset S1**).

A comparison of the overall functions captured by iNgo_557 compared to Nmb_iTM560 and CarveMe automated models, as assessed by KEGG reaction categories, is presented in **Fig 1B**. The overall quality of the reconstruction was assessed using MEMOTE (37). The cumulative MEMOTE score of iNgo_557 was 91% (**Fig 1C**).

### Validation of predictions in iNgo_557 with experimental phenotypes

*In silico* predictions of biomass flux and amino acid supplementation for iNgo_557 were performed and compared to experimental data to validate the model. First, the compositions of three media used for Gc growth were determined: Gonococcal Base Liquid (GCBL), Morse’s Defined Media (MDM), and Roswell Park Memorial Institute 1640 media (RPMI). The metabolites present in each media were assigned to corresponding model exchanges in equivalent amounts and deemed “equally-scaled” media (**Dataset S2**). These simulated media were used to compute biomass flux and consequent predictions of Gc doubling time. Doubling time predictions made with iNgo_557 were then compared to experimental values by conducting growth curves of FA1090 Gc in each of these media (**Fig 2A and B**). The bacterial doubling times predicted by iNgo_557 for equally-scaled media were within 13, 15, and 34 minutes of experimentally determined values in GCBL, MDM, and RPMI, respectively (**Fig 2C**). All predicted growth rates were faster than what was measured experimentally, which is consistent with the structuring of metabolic network models to predict optimal growth (38).

**Fig 2.**
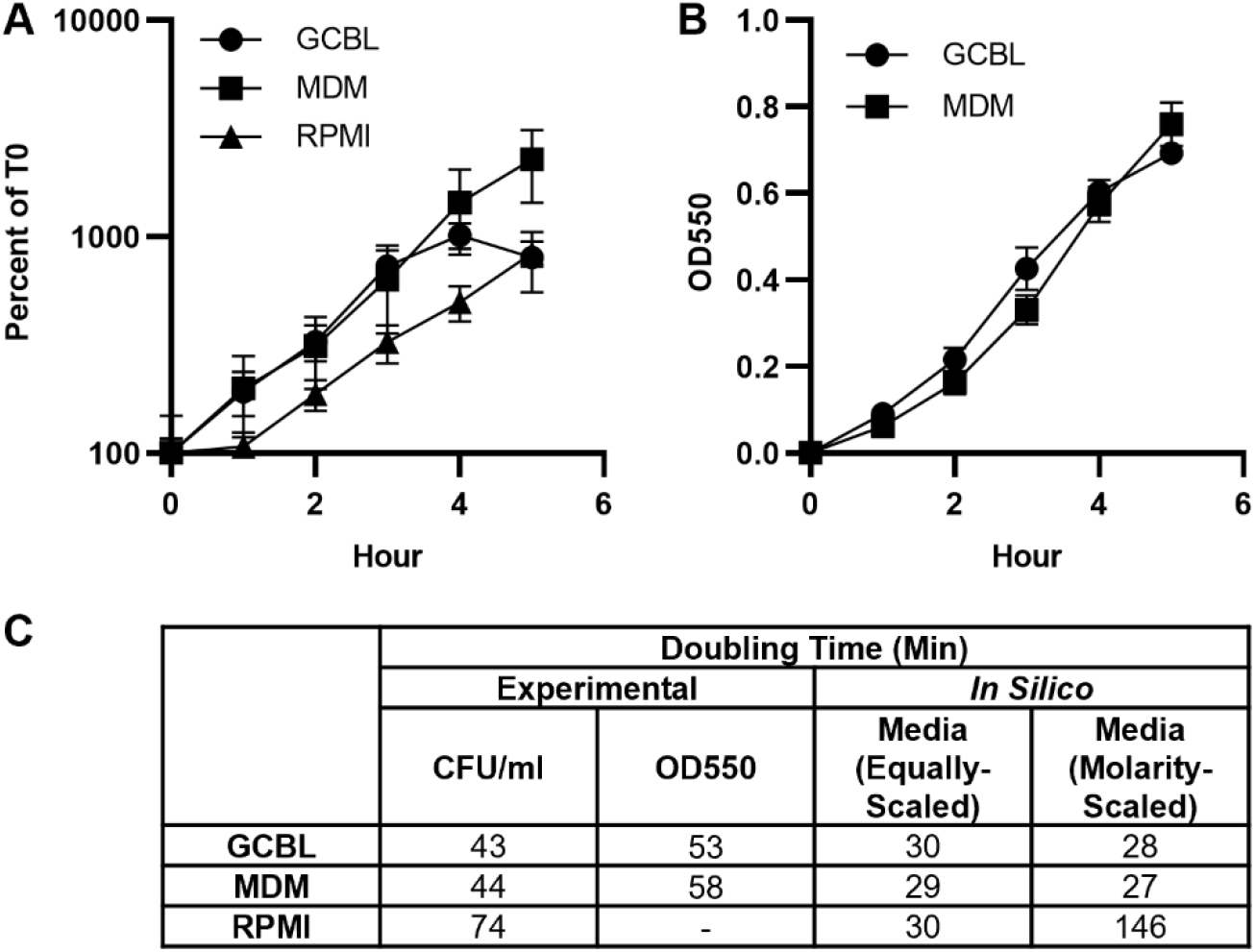
iNgo_557 predicts doubling times that reflect relative growth of Gc in three culture media. Log phase WT Gc was backdiluted into GCBL, MDM, or RPMI and grown over 5 hours. Growth was monitored by (A) enumeration of CFU/ml, reported as percent of CFU measured at 0 hours or (B) optical density at 550 nm. Optical density for Gc grown in RPMI was not determined due to the presence of phenol red indicator that interfered with the readings. n = 4-5 biological replicates. Symbols represent the mean. Error bars represent SEM. (C) Doubling time from A and B was calculated for Gc grown in each medium using GrowthCurver and compared to the predicted doubling times *in silico* using the equivalent concentrations of each nutrient in the different media.

As shown in **Fig 2A**, growth on RPMI was the slowest experimentally, reflecting the limited nutrient content in this media relative to MDM and GCBL (**Dataset S2**). Specifically, metabolite concentrations in RPMI are ~2 to 10-fold less than the concentrations in MDM and GCBL. For example, glucose is found in MDM and GCBL at 27.8 and 22.2 mM respectively, but in RPMI at 11.1 mM (**Dataset S2**). Based on these differences, these three media were molarity-scaled for simulation in iNgo_557, which sets exchanges to be equal to the molarity of each respective metabolite, as has been done previously (39) (**Fig 2C**). While using molarity-scaled media for the substrate concentrations did not change growth predictions for Gc in MDM and in GCBL, the predicted doubling time of Gc in RPMI was substantially slowed, from 30 to 146 min (**Fig 2C**).

To identify the substrate(s) that were limiting for Gc growth in RPMI compared with MDM or GCBL, we used iNgo_557 to predict the growth rate of Gc in a revised formulation of RPMI that was supplemented with 5X the concentration of each component in the original medium (**Table S1**). In simulation of growth in equally-scaled RPMI, only glucose and serine were predicted to increase Gc growth rate, while in the simulation of molarity-scaled RPMI, serine, asparagine, proline, aspartate, glutamate, and glycine were predicted to increase Gc growth rate (**Fig 3A, Table S2**). We tested these predictions experimentally. Addition of 5X glucose, serine, or asparagine to RPMI significantly increased the growth rate of Gc compared with unmodified RPMI (**Fig 3B**). Addition of proline, aspartate, glutamate, or glycine exhibited a trend towards increased growth, though growth differences from unmodified RPMI were not statistically significant. As negative controls, the experimental growth rate of Gc was unaffected when RPMI was supplemented with 5X threonine or valine, which were not predicted to increase growth (**Fig 3B**). These findings demonstrate that iNgo_557 can accurately predict those nutrients that stimulate Gc growth.

**Fig 3.**
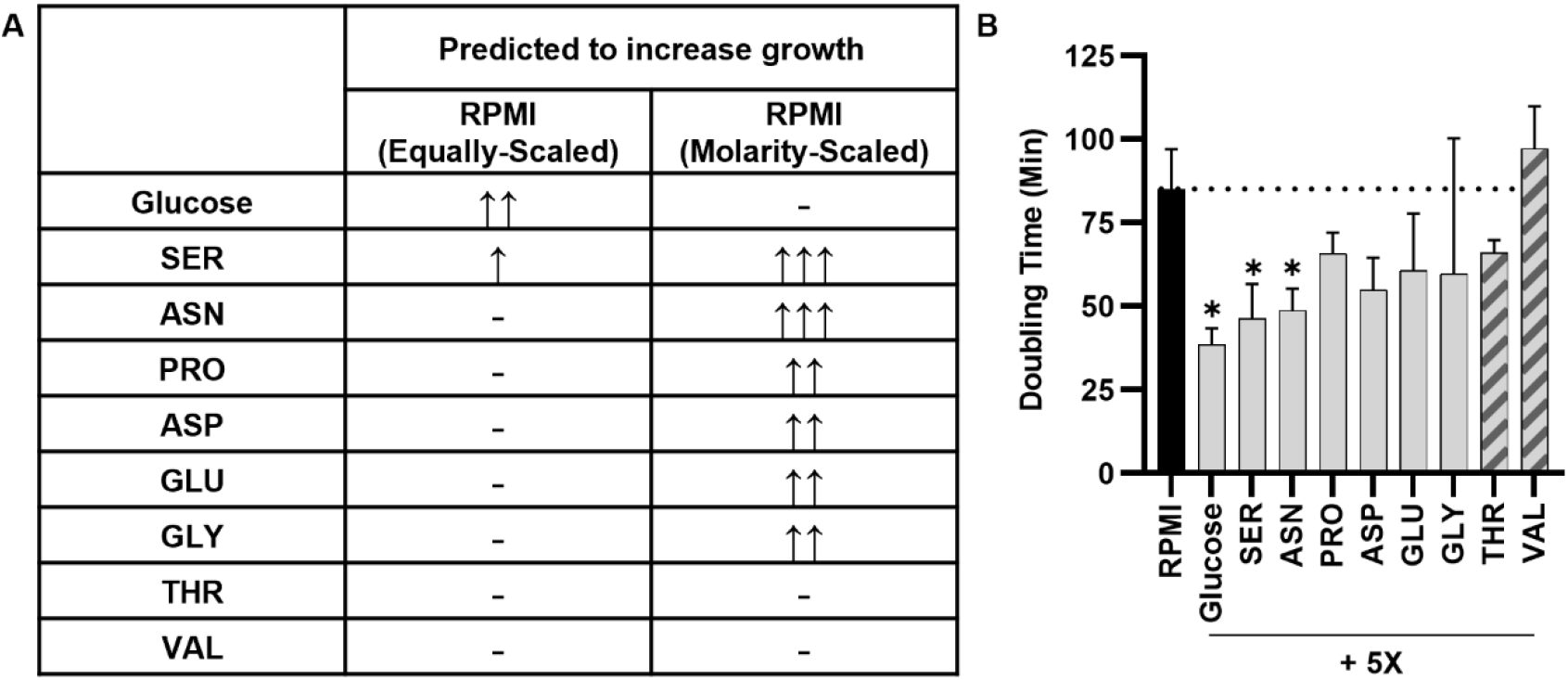
Identification of nutrients that limit Gc growth in RPMI. A) Metabolites in RPMI that are predicted using iNgo_557 to increase Gc growth when increased by 5X the standard flux. Increase in doubling time represented by ↑ ≥ 10%, ↑↑ ≥ 20%, and ↑↑↑ ≥ 30%. B) WT Gc was grown in RPMI supplemented with 5X the concentration of the indicated metabolites for 5 hours. Growth was monitored by enumeration of CFU/ml, and doubling time was calculated using GrowthCurver. Results are from n = 3 biological replicates. Bars represent the mean. Error bars represent SEM. Dotted line indicates doubling time in unmodified RPMI (black bar). Metabolites predicted to increase growth are in gray bars; control metabolites predicted to not increase growth are in hatched bars. *, *P* < 0.05 by one-tailed *t* test relative to unmodified RPMI.

Gc is reported to be capable of growing on only three carbon sources: glucose, lactate, and pyruvate (3). While iNgo_557 successfully predicted growth of Gc on glucose, lactate, and pyruvate as carbon sources in molarity-scaled MDM (27, 43, 28 min doubling times), it also predicted slow growth of Gc in their absence (247 min doubling time). Gc possesses pathways for catabolism of amino acids, which in some other bacteria serve as carbon sources. However, Gc was unable to grow in MDM that did not have one of these carbon sources added (**Fig S3A**), even with additional amino acids experimentally added (**Fig S3B**). Despite this discrepancy between predicted and experimental growth, amino acid catabolic pathways were left intact, to account for Gc usage of amino acids in the presence of its known carbon sources.

One metric commonly used for model validation is the comparison of gene essentiality predictions generated with metabolic reconstructions with those that are identified as essential through transposon mutagenesis (40). We compared gene essentiality predictions yielded by iNgo_557 on GCB to a published dataset that is comprised of essential genes, which were identified through the growth of strain MS11 transposon insertion mutants on GC agar (**Dataset S3**) (41). In iNgo_557, a gene was predicted to be essential if less than 10% of the optimal biomass of the WT could be produced by a mutant in single-gene deletion simulations. Gene essentiality was predicted with an accuracy of 73% and a Mathews Correlation Coefficient (MCC) of 0.43 (**Dataset S3**). Genes correctly identified as essential included those related to LOS and peptidoglycan biosynthesis, purine metabolism, and pyruvate metabolism. Of the genes identified as non-essential by iNgo_557 but essential by transposon library growth assays, many encoded participants in pyrimidine metabolism, oxidative phosphorylation, and glycolysis. We verified that one of these genes, encoding pyruvate kinase (*pyk*), could be deleted from Gc and that the resulting null mutant could grow in GCBL containing glucose as the sole carbon source, albeit slower than the WT parent or when pyruvate was provided (**Fig S4**). This discrepancy between predicted and experimental results could be due to a number of issues, including the fact that these genes could be essential for growth in a competitive setting when mixed with a library of other transposon mutants, differences in media composition between GC agar and GCBL, or differences in Gc strain background (42, 43). As such, arguments for a more nuanced use of gene essentiality data to validate model predictions have been previously made (42). For these reasons, we did not use gene essentiality data for further curation of the reconstruction, but they are included here for reference.

### Transcriptome-guided modeling of Gc metabolism during co-culture with primary human neutrophils predicts a shift in the pyruvate axis

GENREs serve as a tool for scaffolding complex metabolic information in human-interpretable formats. One such application is the integration of transcriptional data with GENREs to develop a comprehensive picture of bacterial metabolism in complex and uncharacterized environments (44). Given that Gc is a human-specific pathogen, we sought to use the reconstruction to predict metabolic phenotypes that are consistent with Gc growth in the context of human neutrophils (PMNs), the predominant immune cell that is recruited during infection. To investigate how Gc metabolism shifts in response to co-culture with PMNs, transcriptomic data from Gc co-incubated with PMNS for 1 hour was integrated with iNgo_557 to generate contextualized models that offer insight to the metabolic state of Gc during infection.

To accomplish this goal, we applied the RIPTiDe (Reaction Inclusion by Parsimony and Transcript Distribution) algorithm (45), which uses RNA-seq data to identify the most cost-effective usage of metabolism while also reflecting the organism’s transcriptional investment. RIPTiDE has been used successfully with models of *Pseudomonas aeruginosa* and *Clostridioides difficile* to uncover metabolic contributors to virulence in the context of mucin degradation, biofilm formation, murine infection models, and co-culture with other microbes (19, 21, 46). We reasoned this approach would generate context-specific models of the metabolism of Gc when grown with and without PMN co-culture and would identify those reactions that are likely to be differentially active in each condition. The transcriptome data set we used was from a constitutively opacity protein-deficient isolate of strain FA1090 Gc, which was cultured in RPMI + 10% fetal bovine serum for 1 hour. Gc was cultured in the presence or absence of primary human PMNs that were adherent and treated with the chemokine interleukin-8 to reflect the activated state of immune cells during infection (47). This isolate of Gc remains primarily extracellular when exposed to PMNs (48). RIPTiDe generated two context-specific models of Gc metabolism: one for Gc in medium without PMNs, and one for Gc with PMNs. For each of the two models, flux samples were generated to assess all possible metabolic profiles in the two environmental contexts. Flux samples generated with the models significantly correlated with the transcript abundances derived from RNAseq for each condition (r=0.242, p<0.001 for Gc without, and r=0.263, p<0.001 for Gc with PMNs), indicating that the context-specific metabolic profiles predicted with RIPTiDe align with experimental data.

Biomass flux was significantly increased in the contextualized model of Gc co-cultured with PMNs, compared with Gc cultured without PMNs (**Fig 4A**), suggesting an overall stimulation of Gc metabolism in the presence of PMNs. Flux distributions for each model were then compared using non-metric multidimensional scaling (NMDS) of consensus reactions shared between both models to broadly identify metabolic growth patterns used by the context-specific models. NMDS revealed that the sampled flux distribution for Gc co-cultured with PMNs overlapped with, but was distinct from, the sampled flux distribution for Gc cultured without PMNs (**Fig 4B**). This result reflects that the media used for growth is consistent between the two models, but co-culture with PMNs caused a shift in metabolic pathways used for growth.

**Fig 4.**
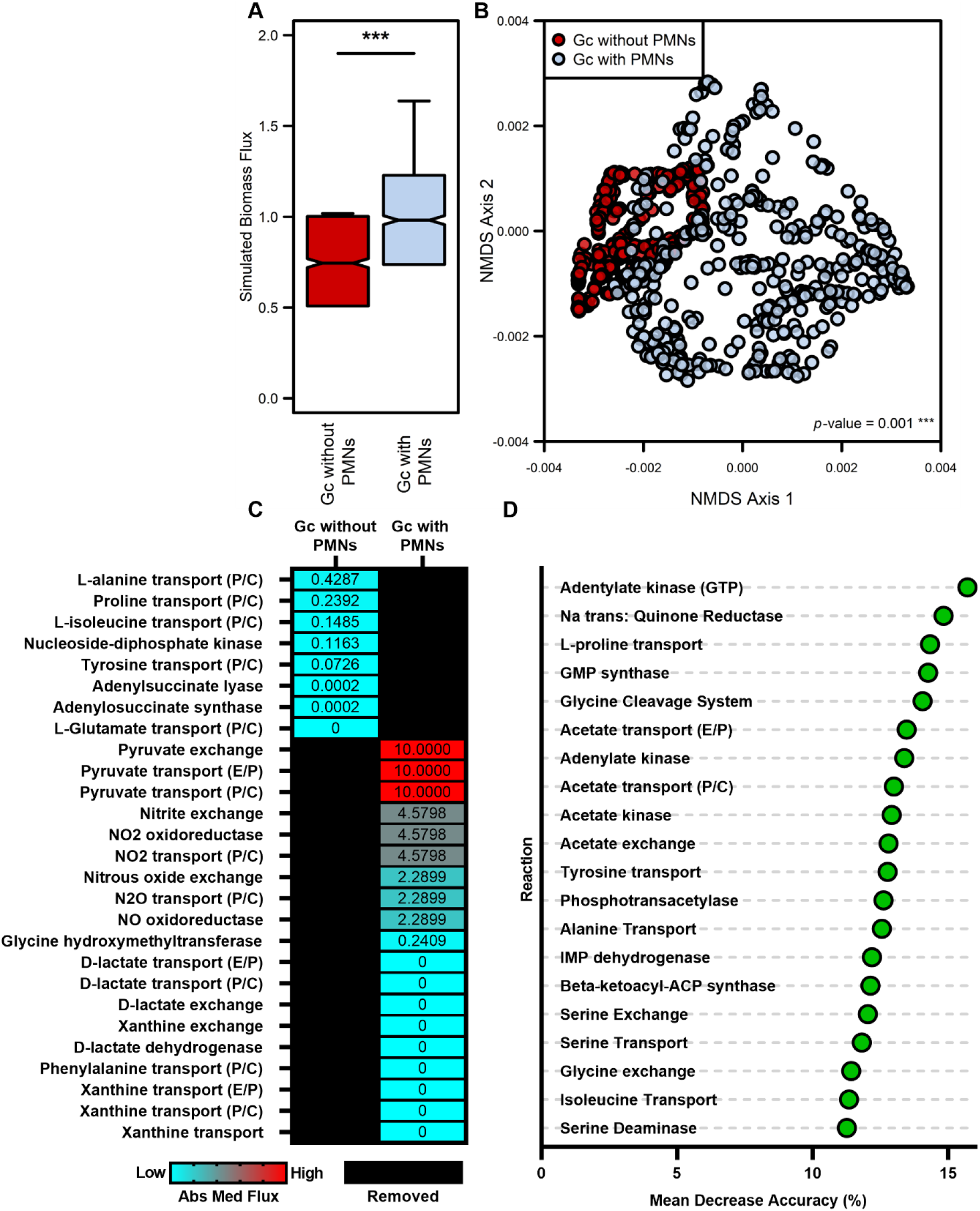
Metabolic activity predictions differ between Gc cultured without PMNs and Gc co-cultured with PMNs. Transcriptomes from Gc cultured with and without PMNs for 1 hour were used to generate context-specific models of iNgo_557 using RIPTiDe. Inactive reactions were pruned during contextualization. (A) Boxplot of biomass objective flux distributions (n=500) from each context-specific model. Significance determined by Wilcoxon rank sum test (*P* value < 0.001). (B) Axes 1 and 2 of four-dimensional Bray-Curtis based NMDS ordination for flux sampling results from non-biomass reactions shared between context-specific models of iNgo_557. Significant difference determined by PERMANOVA. (C) The median absolute value of reaction activities for uniquely active metabolic reactions in each context-specific model. Black boxes indicate reactions are absent in the corresponding model. (D) Random Forest supervised machine learning was used to categorize flux sample activity as Gc without PMNs and Gc with PMNs for non-biomass metabolic reactions shared between the contextualized models. The mean decrease accuracy, which predicts the impact of removal of the reaction from the model on Random Forest categorization predictions (Gc without PMNs vs Gc with PMNs), for the top 20 most differentiating reactions is shown.

We further analyzed the contextualized models to better understand the shifts in metabolism that resulted in the distinctions observed in the NMDS analysis. Reactions unique to each model (non-consensus reactions) were identified, and the absolute median activity for each reaction was determined to examine the contribution of each reaction to biomass production (**Fig 4C**). From this analysis, we identified a set of 19 reactions that were unique to Gc co-cultured with PMNs and 8 reactions unique to Gc cultured without PMNs. Several reactions involved in metabolite import and catabolism were unique to Gc co-cultured with PMNs, suggesting that there are changes to the metabolites available to Gc in this condition, possibly due to competition with or excretion by PMNs. Specifically, pyruvate and D-lactate exchange reactions were unique to Gc co-cultured with PMNs, suggesting bacterial use of these alternative carbon sources in this infection condition (**Fig 4C**). This observation aligns with extensive evidence that PMNs secrete lactate as a byproduct of oxidative metabolism, which stimulates Gc growth (5, 49). Similarly, Gc co-cultured with PMNs were also predicted to uniquely carry flux through nitrogen metabolism, in particular the import of nitric oxide and nitric oxide reductases (**Fig 4C**). These findings align with the reported production of nitric oxide (NO) via inducible nitric oxide synthase (iNOS) in stimulated PMNs (50). Although NO is used by phagocytes to directly kill pathogens, Gc can exploit this aspect of inflammation by detoxifying NO to nitrite, or using nitrite and nitric oxide as terminal electron acceptors during anerobic growth (51). Together, these observations support the hypothesis that neutrophil byproducts mediate remodeling of Gc metabolism.

We next assessed reactions that were shared between both models of Gc cultured without PMNs and Gc co-cultured with PMNs but carried different levels of flux. From this, we identified reactions that most strongly discriminated between metabolic activity of the two models. This analysis employed a supervised machine learning approach with Random Forest, a categorization algorithm that can segregate flux samples based on the contextualized models (**Fig 4D**). We then assessed mean decrease accuracy (MDA) to identify reactions that, when removed from the model, most affected the categorization predictions of the Random Forest. Gc grown in the presence and absence of PMNs were particularly distinguished by flux out of the pyruvate node, through acetate synthesis. Specifically, acetate exchange, acetate transport, acetate kinase, and acetate phosphotransacetylase were identified as reactions that impacted the categorization capabilities of the Random Forest (MDA ~13%) (**Fig 4D**). Acetate production is a prominent feature of bacterial overflow metabolism, in which ATP is generated from the production of acetate from acetyl-CoA via the PTA-AckA pathway rather than shuttled into carbon backbones for biomass (52). Visualization of flux balance analysis (**Fig 5A and B**) demonstrated a predicted increase in acetate flux in co-culture with PMNs, consistent with increased carbon flux from the addition of alternative carbon sources, such as lactate and pyruvate.

**Fig 5:**
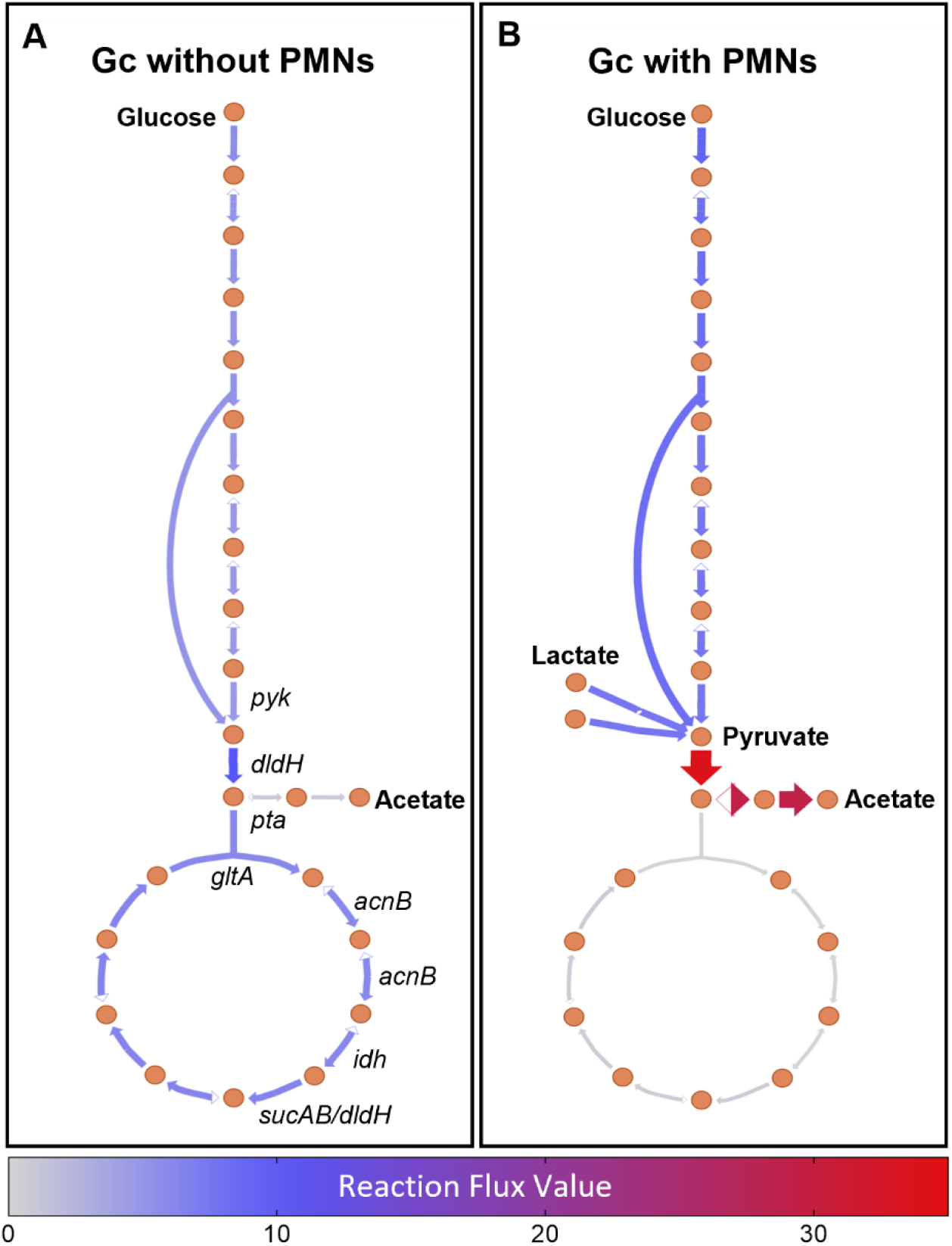
Visualization of flux balance analysis for central carbon metabolism in contextualized models of Gc with and without PMNs. Orange circles indicate metabolites. Relevant imported and exported metabolites are indicated in bold. Arrows indicate reactions. The intensity of coloration and the arrow size indicate the degree of flux through reactions. Conditionally essential genes corresponding to reactions are indicated in italics. Schematics were generated with Escher.

Conditionally essential genes were predicted by conducting essential gene calculations in each model, then comparing between them (**Table 1**, **Dataset S4**). Twelve genes were predicted to be essential only when Gc was cultured without PMNs, and 2 genes were predicted to be essential only when Gc was co-cultured with PMNs. Of the 12 genes predicted to be essential only when Gc was cultured without PMNs, 7 are within a single pathway exiting the pyruvate synthesis node (**Table 1**): pyruvate kinase (*pyk*), portions of the pyruvate dehydrogenase complex (*dldH*), phosphate acetyltransferase (*pta*), citrate synthase (*gltA*), aconitase (*acnB*), and isocitrate dehydrogenase (*idh*) were all predicted to be essential only for Gc cultured without PMNs.

**Table 1:** Conditionally essential genes predicted by single gene deletion analysis of contextualized models of Gc without PMNs and Gc with PMNs.

| Ngo ID | Annotation | Gene ID |
| --- | --- | --- |
| *Gc co-cultured with PMNs* |  |  |
| NGO0799 | inosine-5-monophosphate dehydrogenase | *imdH* |
| NGO2164 | GMP synthase | *guaA* |
| *Gc cultured without PMNs* |  |  |
| NGO0214 | phosphate acetyltransferase | *pta* |
| NGO0562 | dihydrolipoamide dehydrogenase | *dldH* |
| NGO0918 | citrate synthase | *gltA* |
| NGO0925 | dihydrolipoamide dehydrogenase | *dldH* |
| NGO1082 | isocitrate dehydrogenase | *ldh* |
| NGO1231 | aconitate hydratase | *acnB* |
| NGO1325 | glycine dehydrogenase | *gcvP* |
| NGO1404 | glycine cleavage system protein H | *gcvH* |
| NGO1406 | glycine cleavage system protein T | *gcvT* |
| NGO1470 | NAD(P) transhydrogenase subunit alpha | *pntA* |
| NGO1472 | NAD(P) transhydrogenase subunit beta | *pntB* |
| NGO1881 | pyruvate kinase | *pyk* |

To test the prediction that pyruvate synthesis genes were essential for Gc in rich growth medium but dispensable for Gc in the presence of PMNs, we generated a null mutant in pyruvate kinase (Δ*pyk*), the first enzyme in this pathway. As expected, *Δpyk* had a growth defect in MDM containing glucose as the sole carbon source, while the WT parent grew in this medium (**Fig 6A**). Also as expected, *Δpyk* and WT Gc grew equally well in MDM containing either lactate or pyruvate as the sole carbon source (**Fig 6B and C**). We then measured growth of WT and *Δpyk* Gc in the conditions used to collect the PMN transcriptomics data. In RPMI + 10% FBS, the *Δpyk* mutant stopped growing after 3 hours, and by 24 hours its viability had declined to 1 % of the inoculum. In contrast, when cultured in the presence of PMNs, *Δpyk* Gc grew significantly better than Gc in the absence of PMNs, and in fact increased in viability over 24 hours (**Fig 6E, F, G, and H**). WT Gc grew over this time whether or not PMNs were present (**Fig 6D, F, G, and H**). These results suggest that Gc co-cultured with PMNs has a decreased need for flux through glycolysis and instead imply that Gc has access to alternative carbon sources such as lactate and pyruvate, which support its growth in the presence of PMNs independently of the glycolytic pathway.

**Fig 6.**
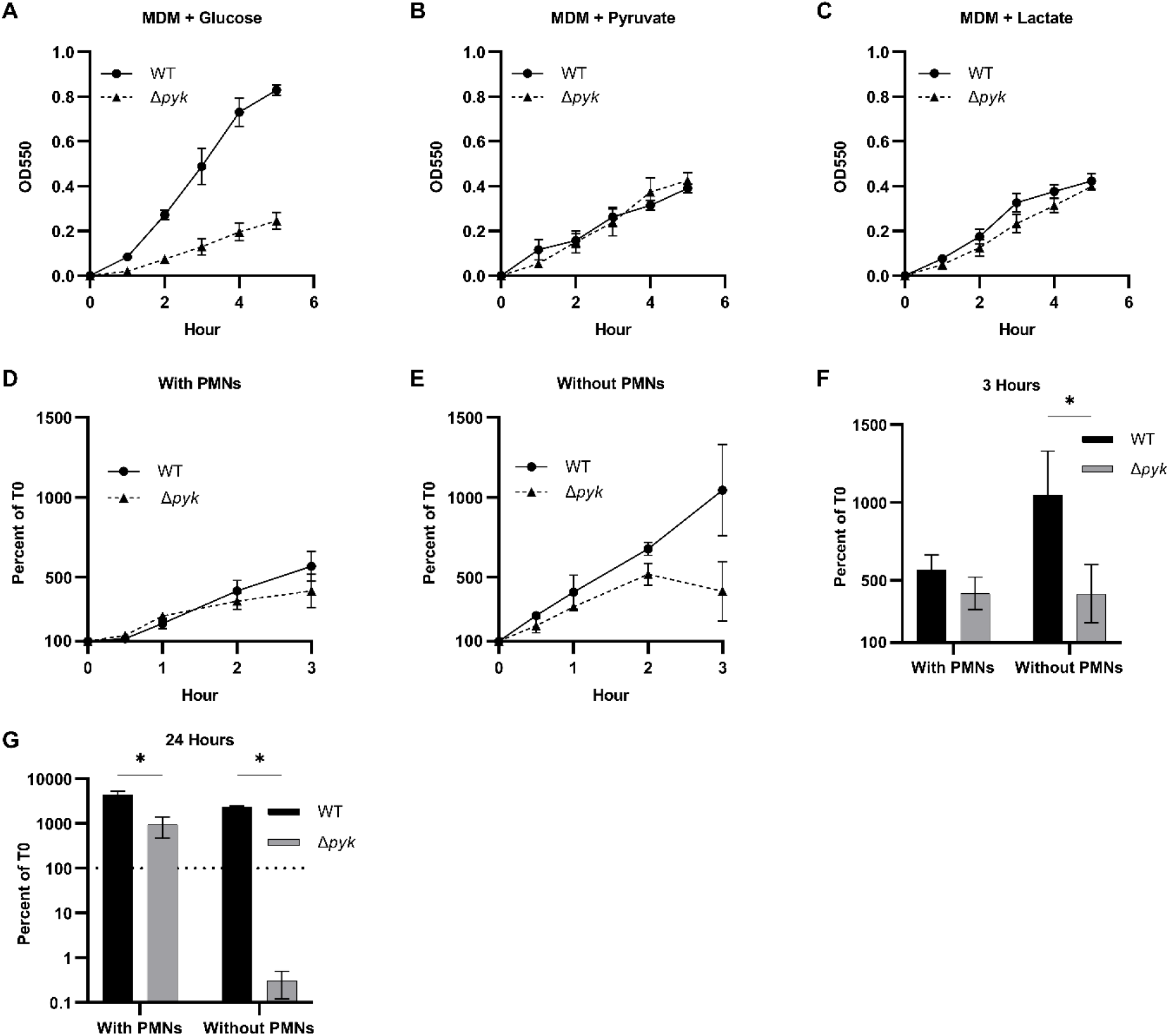
Pyruvate kinase is conditionally essential for *N. gonorrhoeae* in glucose-containing medium, but not for bacteria cultured with PMNs. WT Gc and isogenic Δ*pyk* mutant were cultured in MDM containing (A) glucose, (B) pyruvate, or (C) L-lactate as the primary carbon source. Growth over 5 hours was monitored by optical density at 550 nm for n = 3 biological replicates. Symbols represent the mean. Error bars represent SEM. (D-G) WT and *Δpyk* Gc were exposed to primary human PMNs in suspension or inoculated in RPMI + 10% FBS. CFU were enumerated at 0.5, 1,2, 3, and 24 hours, and Gc growth is reported relative to CFU for that strain at 0 hour (100%). (D and E) Growth curves with (D) and without (E) PMNs over 3 hours. (F and G) Gc CFU at (F) 3 and (G) 24 hours, reported as the percent of CFU for that strain at 0 hours (100%). Bars represent the mean. Error bars represent SEM. n=3 biological replicates. Significance was determined by two-way ANOVA with Holm–Sidak correction for multiple comparisons, * p < 0.05.

## Discussion

Over the last twenty years, genome-scale metabolic modeling has become a powerful tool for context-specific interrogation of complex biological networks. In this study, we developed a highly curated genome-scale metabolic network reconstruction, titled iNgo_557, for Gc strain FA1090. This model predicts the use of glucose, lactate, and pyruvate as carbon sources for Gc, and an increase in growth when selected amino acids are supplemented in cell culture medium containing one of these carbon sources (3). iNgo_557 was contextualized using transcriptomics data that we recently generated (47) to identify shifts in Gc metabolism that occur in response to co-culture with PMNs. These results represent the first use of genome-scale metabolic modeling in Gc for discovery of metabolic contributors to virulence.

Through the linkage of gene, reaction, and metabolite information, iNgo_557 facilitates rapid and convenient manipulation of metabolic parameters to identify contributors towards Gc pathogenesis that are otherwise complicated, time-consuming, or laborious to replicate *in vitro*. Independently, GENREs can be used to simulate well defined environmental contexts, such as growth in laboratory media. We developed *in silico* representations of three commonly used media for Gc. iNgo_557 accurately reflects experimental growth phenotypes in these media and can be used to predict Gc growth phenotypes following distinct manipulations to these media. We demonstrated one such use: identification of growth-limiting nutrients in RPMI. Other applications include nutrient drop-out experiments, aerobic and anaerobic growth, and gene essentiality studies.

While the predictions generated by our model were consistent with experimental results, incorrect predictions are also informative, revealing points of obscurity in our understanding of Gc metabolism. For example, iNgo_557 predicted growth of Gc in MDM in the absence of a dedicated carbon source (glucose, lactate, pyruvate). Upon further interrogation, the predicted growth of Gc on MDM without a carbon source was due to consumption of serine and alanine as carbon sources. Although Gc encodes the genes necessary to catabolize these amino acids (ALATA_L/ NGO_1047 and SERD_L/NGO_1773 and NGO_0444), it is unable to use amino acids as a sole carbon source (**Fig S3**). In other bacteria, such as *P. aeruginosa*, transcriptional and post-transcriptional regulation of serine catabolism has been found to prevent the use of serine as a sole carbon source (53). Our results suggest that a similar form of transcriptional regulation may also dictate Gc carbon source utilization. These discrepancies serve as points for further investigation and facilitate hypothesis generation.

Incorporation of additional layers of regulatory information can improve model accuracy, particularly for the modeling of complex environments such as during co-culture with other species or cell types, which is impeded by lack of knowledge of the metabolite environment. As a human-adapted mucosal pathogen, Gc must co-exist with a complex assortment of human microbiota, epithelial cells, and mucosal immune cells. The recruitment of PMNs and the inflammation associated with gonococcal infection further complicate an already complex metabolic environment. Transcriptomic integration with metabolic models serves to deconvolute the modeling of these complex settings through unsupervised contextualization of GENREs for a specific environment.

As such, we leveraged RIPTiDe with iNgo_557 to better understand the metabolic pathways enabling Gc growth during co-culture with PMNs and to predict the behaviors of this host-associated bacterial species. Intriguingly, several genes are predicted to be essential only when Gc is cultured without PMNs, but not in the context of PMNs. The majority of genes predicted to be essential in the absence of PMNs were downstream of pyruvate kinase (*pyk*) within pathways exiting the pyruvate synthesis node. We validated this prediction by showing that Gc required pyruvate kinase for growth in rich medium, but not when co-cultured with PMNs. Metabolic modeling using iNgo_557 predicts that this effect is due to the bypass of pyruvate synthesis through import of alternative carbon sources, including lactate and pyruvate, when in the presence of PMNs, which is supported by our growth data. Our results align with previous reports showing the ability of Gc to consume lactate and pyruvate derived from host cells (5, 49, 54). PMNs are highly glycolytic cells, consuming glucose and secreting lactate following stimulation with PAMPs (49). Use of lactate was previously reported to be required for Gc survival from PMNs, within cervical epithelial cells, and in the female mouse genital tract (5, 6, 55). The increase in biomass flux predicted for models of Gc cultured with PMNs compared to Gc cultured without PMNs is further consistent with reports that Gc growth on lactate stimulates Gc metabolism (49, 55). Together these results provide evidence that Gc utilizes addition alternative carbon sources, such as lactate and pyruvate, when co-cultured with PMNs to enhance its growth.

Regardless of the source, carbon exiting the pyruvate synthesis node, can proceed in one of two pathways in Gc: acetate production or oxidation through the TCA cycle. Acetate production through the PTA-AckA pathway is a prominent feature of Gc growth on glucose, lactate, and pyruvate (52, 56). Downstream of pyruvate kinase, iNgo_557 predicted increases in Gc acetate production when in the presence of PMNs. In *N. meningitidis*, acetate is secreted following growth on glucose, lactate, and pyruvate, and the highest activity of the PTA-AckA pathway occurs when all three carbon sources are present, compared with glucose alone (56). Our results are consistent with this observation. Alternatively, glucose, lactate, and pyruvate can instead be further catabolized by the TCA cycle. In *N. meningitidis*, pyruvate dehydrogenase (*dldH*), citrate synthase (*gltA*), aconitase (*acnB*), and isocitrate dehydrogenase (*idh*) reaction activities were all demonstrated to be high in the presence of glucose, but decreased in the presence of pyruvate (56). Consistent with the stimulation of these enzymes in the presence of glucose compared to pyruvate, iNgo_557 predicted *dldH, gltA, acnB*, and *idh* to be essential only in the absence of PMNs, in which glucose is the sole carbon source available. The alleviation of the requirement for *acnB* in the context of PMN co-culture is notable in light of a recent study that identified compensatory mutations within *acnB* that enabled the recovery of antibiotic-resistant *penA* mutant Gc from the mouse genital tract (57). Together our results highlight the pyruvate node as a critical pivot point in Gc metabolism, particularly in the context of an inflammatory environment created by PMNs. Overall, the predictions generated here by contextualized models of iNgo_557 reveal new insights into Gc pathogenesis, highlighting it as a viable platform for the discovery of metabolic pathways associated with virulence and antibiotic resistance.

Treatment options for Gc have become increasingly limited over the last two decades, and only a single recommended antibiotic remains for the treatment of gonorrhea (58). The development of new potential therapies is essential to avoid the threat of completely antibiotic-resistant Gc. Targeting essential bacterial metabolic pathways during infection represents a promising approach, one that was first shown decades ago in the context of sulfonamide antibiotics, which directly inhibit folate synthesis (59). Novel approaches for the treatment of antibiotic-resistant infections have included the application of metabolites to shift the metabolism of pathogens towards a less favorable state (60). There is a need for a revisitation of Gc metabolism and physiology in light of the approaching post-antibiotic era for gonorrhea (61). Technologies such as RNA-sequencing, forward and reverse genetic screens, and metabolic modeling can all provide insights into Gc metabolism. Here, the integration of transcriptomics with genome-scale metabolic modeling is synergistic, providing more insight into the remodeling of Gc metabolism in the context of PMN co-culture than could be discerned from each technique alone. In sum, this study highlights the opportunities afforded by genome-scale metabolic modeling for targeted identification of context-specific essential metabolic pathways that enable Gc to thrive within the human host, with further predictions and discoveries remaining to be made.

## Methods

### Genome-scale metabolic reconstruction

To generate a GENRE for Gc, we used *N. meningitidis* M58 Nmb_iTM560 as an initial template for the automated multi-strain model, reconstruction pipeline (26). In brief, the pipeline used bidirectional best hit BLAST to identify genes with >80% homology between *N. meningitidis* M58 (AE002098.2) and *N. gonorrhoeae* FA1090 (AE004969.1) to generate a homology matrix for the two species. A secondary comparison using BLAST on nucleotide sequences was conducted to identify potential homologs with poor ORF annotation. These automated calls were inspected and reassessed for each gene present in Nmb_iTM560 as indicated in **Dataset S1**. Using the homology matrix, a draft strain-specific model was generated using COBRApy (62). Metabolic genes (and the corresponding reactions and metabolites) specific to Gc FA1090 were added to the reconstruction using CarveMe when supported by literature evidence (36). Exchange reactions that were missing for extracellular metabolites in the reconstruction were added. The model was then further manually curated to de-orphan reactions and incorporate published metabolic functions for Gc according to literature evidence where possible (**Dataset S1**). Final gene and reaction calls, along with decision annotations, can be found in **Dataset S1**. Annotation data was automatically assigned using ModelPolisher (63). Reaction and stoichiometric inconsistencies were corrected for each reaction. All formulas were mass and charge balanced using the BIGG database, when possible, in order to maintain a consistent namespace (25). A list of mass and charge imbalanced reactions and their corrections are provided in **Dataset S1**. Additional annotations were collected and added to the annotation field dictionary for all model components from KEGG, PATRIC, Uniprot, MetaNetX, MetaCyc, PubMLST, or BIGG databases (25, 29–33, 64). The pipeline for development of the reconstruction is available in the GitHub repository associated with this study (https://github.com/aimeepotter/Gc_GENRE_2022).

### Assessing reconstruction quality

Modeling assessments, including flux balance analysis, flux-variability analysis, single gene knock-out analysis, were conducted using COBRApy (62). Model quality was assessed with MEMOTE using a local installation v0.13.0 (37). Gene essentiality predictions were compared to a published dataset of essential genes for growth on solid, rich media for Gc strain MS11 (41), which was aligned to Gc FA1090 by bidirectional best hit BLAST as above. Prediction accuracy was calculated as the number of correct predictions divided by the number of total predictions for genes present in both datasets, and the Matthews correlation coefficient (MCC) was calculated as in (65).

A protocol for defining realistic modeling constraints for *in silico* media was recently described, in which metabolite exchanges are scaled based on the maximum possible usage defined by the concentration of metabolites in mmol/L (39). We therefore generated two *in silico* exchange reaction constraints for each simulated media: equally-scaled, to avoid constraining the model with incorrect assumptions, and molarity-scaled, to match the maximum possible use of metabolites. The concentration of metabolites present in each media and their corresponding assignments to *in silico* media constraints are detailed in **Dataset S2**. Biomass flux and subsequent doubling times for simulated growth in GCBL, MDM, and RPMI were compared to experimental values. Predictions of Gc doubling time were calculated assuming a biomass equation scaled to 1g dry weight of bacteria based on the following formula:

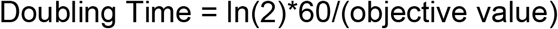

Experimental doubling times were determined using GrowthCurver implemented in R for both OD and CFU/ml, with stationary phase values trimmed (66).

### RIPTiDe (Reaction Inclusion by Parsimony and Transcript Distribution) contextualization & analysis

Transcriptomic data retrieved from the Gene Expression Omnibus (GEO) database (GSE123434) for Gc cultured without and with PMNs over the course of 1 hour was mapped to the corresponding FA1090 gene IDs using the conversion table provided in Dataset S1 of (47). For RIPTiDe contextualization, an unsupervised approach was used in which all exchange reaction bounds were set to ±10, except oxygen, which was set at ±20. The transcriptomic data was then integrated with the model using RIPTiDe using the maxfit_contextualize() function (minimum fraction 0.3, maximum fraction 0.8, n=1000) to produce contextualized models for Gc grown in the presence or absence of PMNs (45). Flux samples were gathered from consensus reactions between both contextualized models (n=500 samples per model). Bray-Curtis based NMDS (k=4, trymax=25) and permutational multivariate analysis of variance (PERMANOVA) (perm=999) analyses were accomplished using the Vegan R package (67). Supervised machine learning was accomplished with the implementation of AUC-Random Forest also in R (68). Statistical analysis was performed in R v4.1.0. Visualizations of flux balance analysis were performed using Escher (69).

### Bacterial strains and growth conditions

Opaless Gc is a non-variable Opa-deficient derivative of the FA1090 background constitutively expressing the pilin variant 1-81-S2, which served as the WT for all experiments (48, 70). Strain 130 Δ*pyk* was generated by transformation with an overlap extension PCR product, replacing the *pyk* ORF with a spectinomycin resistance cassette using the following primers: Pyk Upstream F-CCGAATACGGCGACTTTACC, Pyk-SacI-Omega F-CAAAATCGTCGCCACCCTTGGAGCTCTGCCCGTTCCATACAGAAGC, Pyk upstream R-GCTTCTGTATGGAACGGGCAGAGCTCCAAGGGTGGCGACGATTTTG, Pyk downstream F-GCTCACAGCCAAACTATCAGGTGAGCTCCAGACGGAGTATCCCGAAGC, Pyk-sacI-Omega R-GCTTCGGGATACTCCGTCTGGAGCTCACCTGATAGTTTGGCTGTGAGC, Pyk downstream R-ACTGTGTGCCGAAGTGGTAG. Mutation was confirmed by sequencing and PCR.

WT Gc were grown on Gonococcal Medium Base (GCB, Difco) plus Kellogg’s supplements at 37°C with 5% CO2 (71, 72). *Δpyk* strains were grown on GCB plus Kellogg’s supplements with glucose replaced by pyruvate (36 mM) as in (54). For preparation of mid-logarithmic phase bacteria, Gc were grown in liquid medium (GCBL) or carbon matched GCBL containing pyruvate (45 mM) as the sole carbon source, where appropriate, for successive rounds of dilution, and enriched for piliation, as previously described (73). Spectinomycin was used for selection of the *pyk* mutation at 80 μg/ml.

### Growth Curves

Gc in mid-logarithmic phase were pelleted, resuspended in the indicated media, and diluted to ~5*10^7 CFU/ml in 6 ml of media in 15 ml conical tubes (Sarstedt). The bacterial suspension was incubated with rotation at 37°C. Bacterial growth was measured by OD550 and CFU enumeration at specific timepoints. CFU are presented relative to 0 h (100%). Gc was grown in GCBL, HyClone RPMI 1640 media without glutamine (Catalog#SH30096.FS) (Cytivia), or carbon-matched Morse’s defined media (MDM) containing either glucose (27mM), lactate (54mM) or pyruvate (54mM) (74). Doubling times were calculated from best fit logistic curves generated with GrowthCurver (66) for the lag and exponential phase of each growth curve for at least 3 experimental replicates and averaged. Significant differences for growth over time were determined by one-tailed t-test in Graphpad Prism v9.

### Gc-PMN co-culture

PMNs were isolated from venous blood as previously described and used within 2 h of isolation (73). Subjects gave informed consent in accordance with an approved protocol by the University of Virginia Institutional Review Board for Health Sciences Research (#13909). Synchronized Gc infection of PMNs in suspension was conducted as previously described (75). PMNs were resuspended in RPMI (Cytivia) containing 10% heat-inactivated fetal bovine serum (Gibco) at 1 ×10^6^ PMN/ml and Gc was added to each tube at a multiplicity of infection of 10. Six ml of the suspension was incubated in 15 ml conical tubes with rotation at 37°C. Bacterial CFU were enumerated at specified time points and expressed relative to the CFU at 0 h (100%). Data are expressed as the mean ± SEM of at least three replicate experiments. Significant differences were determined by two-way ANOVA with Holm-Sidak correction for multiple comparisons in Graphpad Prism v9.

## Data and Code Availability

Python and R code/packages/scripts used to perform transcriptomics data analyses and generate figures are available on GitHub at https://github.com/aimeepotter/Gc_GENRE_2022. All RNA-seq data are available in the Gene Expression Omnibus (GEO) database under accession GSE123434 (47).

## Acknowledgements

We thank members of the Papin lab, particularly Matt Jenior and Dawson Payne, for thoughtful feedback on modeling parameters. This work was supported by NIH R01AI097312, NIH R01127793, NIH R21AI161302, and the Harrison Distinguished Teaching Professorship (to AKC). ADP was supported in part by NIH T32AI007496. The authors declare that no competing interests exist.

## Supplemental Material

**Fig S1: Best fit logistic curves generated with GrowthCurver were used to calculate experimental doubling time for Gc grown in GCBL, MDM, and RPMI.** Log phase WT Gc was backdiluted into GCBL, MDM, or RPMI. Growth over 5 hours was monitored by (A) enumeration of CFU/ml or (B) optical density at 550 nm. Optical density for Gc grown in RPMI was not reported due to phenol red indicators in the media. n = 4-5 biological replicates.

**Fig S2: Best fit logistic curves generated with GrowthCurver were used to calculate experimental doubling time of Gc grown in RPMI supplemented with potential limiting metabolites.** Log phase WT Gc was backdiluted into RPMI supplemented with 5X the standard concentration of metabolites indicated in **Fig. 3**. Glucose (GLC), serine (SER), asparagine (ASN), proline (PRO), aspartate (ASP), glutamate (GLU), and glycine (GLY) were predicted to be growth limiting; threonine (THR) and valine (VAL) were not predicted to be growth limiting. Growth over 5 hours was monitored by enumeration of CFU/ml. n = 3 biological replicates per condition.

**Fig S3: *N. gonorrhoeae* requires glucose, pyruvate, or lactate as a carbon source for growth.** Log phase WT Gc was backdiluted into MDM containing (A) no dedicated carbon source (no glucose, lactate, or pyruvate) or (B) with 1% Casamino acids added as the carbon source. Growth was monitored by optical density at 550 nm over 5 hours. (A) n = 3 biological replicates. Symbols represent the mean. Error bars represent SEM. (B) n = 1 biological replicate.

**Fig S4: Growth dynamics of WT and *Δpyk N. gonorrhoeae* in GCBL with different carbon sources.** Log phase WT Gc and an isogenic *Δpyk* mutant were backdiluted into GCBL containing (A) 22 mM glucose or (B) 45 mM pyruvate as the primary carbon source and grown for 5 hours. Growth was monitored by optical density at 550 nm for n = 1 biological replicate.

**Fig S5: A *N. gonorrhoeae pyk* mutant does not grow in RPMI containing glucose as sole carbon source.** Log phase WT Gc and an isogenic *Δpyk* mutant were backdiluted into RPMI. Growth over 5 hours was monitored by enumeration of CFU/ml reported as percent of CFU measured at 0 hours (100%). Symbols represent the mean. Error bars represent SEM. n=3 biological replicates.

**Table S1: Concentrations of nutrients predicted to be limiting for Gc growth in RPMI.**

**Table S2: Growth predictions for *N. gonorrhoeae* in RPMI in equally-scaled or molarity-scaled models when selected metabolites are added at five-fold the original concentration.** ^1^Increase in predicted growth rate when the indicated metabolite is increased by five-fold (5x), expressed relative to growth rate in unmodified RPMI.

**Dataset S1:** Annotations on the curation of reactions, metabolites, and genes of iNgo_557.

**Dataset S2:** *In silico* formulations for GCBL, MDM, and RPMI. Simulated media include an “equally-scaled” and a “molarity-scaled” formulation. “Molarity-scaled” formulation based on calculated molarities of metabolites present in GCBL, MDM, and RPMI.

**Dataset S3:** Essential gene predictions for iNgo_557 on simulated GCBL. Genes were deemed essential if knock-out of the gene resulted in <10% of maximal biomass production. Essential gene predictions were compared to experimentally determined essential genes from MS11.

**Dataset S4:** Essential gene predictions with contextualized models of iNgo_557 for Gc without PMNs and Gc with PMNs. Genes were deemed essential if knock-out of the gene resulted in <10% of maximal biomass production.

